# A human high-fidelity DNA polymerase holoenzyme has a wide range of lesion bypass activities

**DOI:** 10.1101/2024.10.14.618244

**Authors:** Rachel L. Dannenberg, Joseph A. Cardina, Helen Washington, Shijun Gao, Marc M. Greenberg, Mark Hedglin

## Abstract

During replication, lagging strand lesions are initially encountered by high-fidelity DNA polymerase (pol) holoenzymes comprised of pol δ and the PCNA sliding clamp. To proceed unhindered, pol δ holoenzymes must bypass lesions without stalling. This entails dNMP incorporation opposite the lesion (insertion) and the 5’ template nucleotide (extension). Historically, it was viewed that high-fidelity pol holoenzymes stall upon encountering lesions, activating DNA damage tolerance pathways that are ultimately responsible for lesion bypass. Our recent study of 4 prominent lesions revealed that human pol δ holoenzymes support insertion and/or bypass for multiple lesions and the extents of these activities depends on the lesion and pol δ proofreading. In the present study, we expand these analyses to other prominent lesions. Collectively, analyses of 10 lesions from both studies reveal that the insertion and bypass efficiencies of pol δ holoenzymes each span a complete range (0 – 100%). Consequently, the fates of pol δ holoenzymes upon encountering lesions are quite diverse. Furthermore, pol δ proofreading promoted holoenzyme progression at 7 of the 10 lesions and did not deter progression at any. Altogether, the results significantly alter our understanding of the replicative capacity of high-fidelity pol holoenzymes and their functional role(s) in lesion bypass.

## INTRODUCTION

In humans, like all eukaryotes, lagging strand DNA templates are primarily replicated by DNA polymerase δ (pol δ) holoenzymes comprised of pol δ and the PCNA processivity sliding clamp. Pol δ, a member of the highly-conserved B-family of DNA polymerases, contains distinct active sites for stringent DNA polymerase and 3’ → 5’ exonuclease (i.e., proofreading) activities and is often referred to as a “high-fidelity” or “replicative” DNA polymerase (2). PCNA has a central cavity large enough to encircle double-stranded DNA (dsDNA) and slide freely along it (3). Thus, association of pol δ with PCNA effectively tethers the polymerase to DNA, maximizing the efficiency and processivity of DNA synthesis by pol δ.

During scheduled DNA replication in S-phase of the cell cycle, pol δ holoenzymes are the first to encounter lagging strand template nucleotides that have been damaged by covalent modifications. These damaging modifications, referred to as lesions, arise from exposure of genomic DNA to reactive molecules and metabolites as well as environmental toxicants. For replication by a progressing pol δ holoenzyme to proceed unhindered on a damaged lagging strand template, the DNA polymerase must catalyze deoxynucleotide monophosphate (dNMP) incorporation directly opposite the lesion (insertion) and the adjacent template nucleotide (nt) 5’ to the lesion (extension) without dissociating (**Figure 1**). Extension places the 3’-hydroxyl terminus of the elongating primer strand back into a base pair with proper geometry for optimal dNMP incorporation. When both insertion and extension have been completed, the lesion is “bypassed” and the progressing pol δ holoenzyme may continue to catalyze processive dNMP incorporation downstream of the lesion (elongation, **Figure 1**)(1). A lesion to a template nucleotide can alter or eliminate its base pairing properties, obligating a DNA polymerase to generate an incorrect base pair (i.e., mismatch) during insertion. Furthermore, the altered geometry of a mismatch at an insertion site can affect the positioning of the primer’s 3’-hydroxyl terminus and the stacking of a nascent base pair for extension (4).

**Figure 1.**
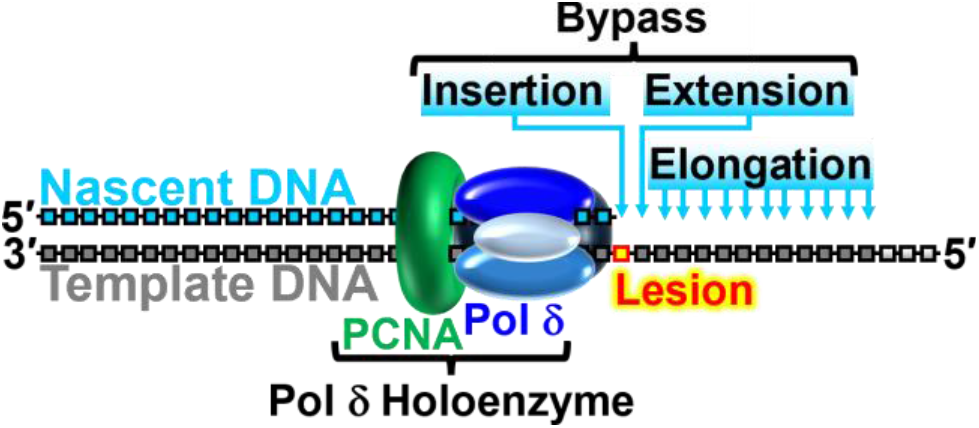
A progressing pol δ holoenzyme encountering a lesion within a lagging strand template.

Considering this and the innate properties of human pol δ discussed above, it had been historically viewed that progressing pol δ holoenzymes stall upon encountering lesions, activating DNA damage tolerance (DDT) pathways that are ultimately responsible for the entirety of lesion bypass. In human cells, the predominant DDT pathway is translesion DNA synthesis (TLS) where one or more specialized TLS DNA polymerases (TLS pols) are recruited to execute lesion bypass. With a more open DNA polymerase active site and the lack of an associated proofreading activity, TLS pols support stable yet potentially erroneous bypass of a diverse array of lesions. This allows processive dNMP incorporation by pol δ holoenzymes to resume downstream of the offending lesion during elongation (5).

In a recent study from our lab (1), we reconstituted human lagging strand replication at physiological pH, ionic strength, and deoxynucleotide triphosphate (dNTP) concentrations to quantitatively analyze, at single nucleotide resolution, the enzymatic activities of pol δ holoenzymes during initial binding events with primer/template (P/T) DNA substrates that each contain a lesion ≥ 9 nt downstream of the P/T junction from which DNA synthesis initiates from. The P/T DNA substrates contained either 7,8-dihydro-8-oxo-2’-deoxyguanosine (8OxoG), 5,6-dihydroxy-5,6-dihydrothymidine (thymidine glycol, Tg), *O*^*6*^-methyl-2’-deoxyguanosine (O6MeG), or *1,N*^*6*^-etheno-2’-deoxyadenosine (εA). Under the conditions of the assay, a given lesion is encountered only by a progressing pol δ holoenzyme rather than pol δ alone. Furthermore, any observed proofreading by pol δ holoenzymes occurs intrinsically (as opposed to extrinsically) because a given dNMP incorporation and subsequent proofreading of that dNMP incorporation are not separated by a pol δ dissociation event (1). The results revealed significant efficiencies of insertion and bypass for multiple lesions. Consequently, stalling of pol δ holoenzymes (via dissociation of pol δ) was distributed among the insertion and extension steps and a portion of the encounters resulted in lesion bypass and elongation to various extents.

Furthermore, proofreading by pol δ holoenzymes did not restrict their progression at any of the aforementioned lesions but rather promoted progression at certain lesions. These results revealed unforeseen complexity and heterogeneity in the replication of lagging strand lesions, challenging our fundamental understanding of lagging strand DNA replication and DDT.

In the present study, we expand these analyses to other prominent lesions to investigate the range of insertion and bypass efficiencies of human pol δ holoenzymes and further decipher the contribution of proofreading to pol δ holoenzyme progression at lesions. A structurally diverse set of lesions produced by a variety of DNA modifying conditions were analyzed. Lesions examined include those resulting from hydrolysis (abasic site, Ab, uracil, U, inosine, I), ultraviolet radiation (cyclobutane thymidine-thymidine dimer T<>T), as well as those produced under oxidative conditions (*N*6-(2-deoxy-α,β-D-erythro-pentofuranosyl)-2,6-diamino-4- hydroxy-5-formamidopyrimidine, FapyG) or by alkylation (3-methyl-2’-deoxycytidine, 3MeC). Collectively, analyses of ten lesions from the previous (1) and present study reveal that insertion and bypass efficiencies of the human pol δ holoenzyme, a high-fidelity DNA polymerase, each span a complete range (from 0 to 100 %). Consequently, the fates of progressing pol δ holoenzymes upon encountering lesions are quite diverse. Furthermore, proofreading by pol δ holoenzymes does not deter their progression at any lesion analyzed. Rather, it significantly promotes progression at 7 of the 10 lesions analyzed. Altogether, the results significantly alter our understanding of the replicative capacity of high-fidelity pol δ holoenzymes and their functional role(s) in lesion bypass on lagging strand templates.

## MATERIAL AND METHODS

### Recombinant Human Proteins

Human RPA, Cy5-PCNA, RFC, and pol δ (exonuclease-deficient and wild-type) were obtained as previously described (6,7). The concentration of active RPA was determined via a FRET-based activity assay as described previously (8).

### Oligonucleotides

All oligonucleotides (except the FapyG-containing 40-mer and control 40-mer described below) were synthesized by Integrated DNA Technologies (Coralville, IA) or Bio-Synthesis (Lewisville, TX) and purified on denaturing polyacrylamide gels. To obtain a template DNA strand containing a FapyG lesion, a 40-mer oligonucleotide containing FapyG was synthesized (5’-ACACAGACGTACTATCATGAC **X**CATCAGACAACGTGCGTC-3’, **X** = FapyG) following an established procedure (9). To ensure an efficient coupling (coupling yield > 90%), the FapyG phosphoramidite concentration was increased to 0.15 M during oligonucleotide synthesis. Purification was carried out via 20% denaturing polyacrylamide gel electrophoresis at ≤ 60 °C. A 40-mer containing G instead of FapyG and TAMRA at the 3’ terminus (but otherwise identical 40-mer FapyG oligo described above) was loaded side-by-side. The ability to see the fluorescently labeled oligonucleotide enables achieving effective separation by running the product close to the bottom of the gel without concern of eluting it from the gel. The fluorescently labeled oligonucleotide (5’-ACACAGACGTACTATCATGACGCATCAGA CAACGTGCGTC-TAMRA-3’) was synthesized with 3’-TAMRA-CPG (Glen Research), deprotected by tBuNH_2_:H_2_O = 1:3 for 6 h at 60 °C, concentrated and used without further purification. The isolated FapyG-containing oligonucleotide was characterized using LC-MS (**Figure S1**). The FapyG-containing 40-mer oligonucleotide was ligated to a 22-mer oligonucleotide (5’-P-AAAAATTACGTGCGGAAGGAGT-3’, P = Phosphate) as follows. Equimolar amounts of the oligonucleotides to be ligated were mixed with a 2-fold molar excess of a 30-mer splint oligonucleotide (5’-CCGCACGTAATTTTTGACGCACGTTGTCTG-3’) in 1X Annealing Buffer (10 mM Tris-HCl, pH 8.0, 100 mM NaCl, 1 mM EDTA), the resultant solution was heated to 60 °C for 5 minutes and allowed to slowly cool to room temperature. T4 DNA Ligase Reaction Buffer (NEB) was added to a 1X final concentration (50 mM Tris-HCl, 10 mM MgCl_2_, 1 mM ATP, 10 mM DTT, pH 7.5). The ligation reaction was initiated by the addition of T4 DNA ligase (NEB), and the resultant solution was incubated for 16 h at 16 °C. Finally, the ligated FapyG-containing 62-mer was purified via 8% denaturing polyacrylamide gel electrophoresis at ≤ 60 °C.

The concentrations of unlabeled DNAs were determined from the absorbance at 260 nm using the calculated extinction coefficients. The concentrations of Cy5-labeled DNAs were determined from the extinction coefficient at 650 nm for Cy5 (ε_650_ = 250,000 M^−1^cm^−1^). The concentrations of Cy3-labeled DNAs were determined from the extinction coefficient at 550 nm for Cy3 (ε_550_ = 125,000 M^−1^cm^−1^). For annealing two single strand DNAs, the 29-mer primer and corresponding complementary 62-mer template strands were mixed in equimolar amounts in 1X Annealing Buffer (10 mM Tris-HCl, pH 8.0, 100 mM NaCl, 1 mM EDTA), heated to 95 °C (60 °C for FapyG-containing 62-mer templates) for 5 minutes, and allowed to slowly cool to room temperature. Due to the instability of a native abasic site during oligonucleotide synthesis and purification, tetrahydrofuran is utilized as an abasic site mimic in which a methylene group replaces the 1 position of 2-deoxyribose (10).

### Steady State Primer Extension Assays

All primer extension experiments were performed at physiological pH (7.5), ionic strength (200 mM), and concentrations of each dNTP exactly as described previously (1). For all experiments, the final ionic strength was adjusted to 200 mM by addition of appropriate amounts of KOAc and samples were protected from light whenever possible. The final concentrations of all reagents, substrate, and protein concentrations were as follows; 1X Replication Buffer (25 mM HEPES, pH 7.5, 10 mM Mg(OAc)_2_, 125 mM KOAc), 1 mM DTT, 1 mM ATP, 250 nM Cy5-labeled P/T DNA (**Figure S2**), 1 μM Neutravidin, 750 nM RPA (heterotrimer), 250 nM PCNA (homotrimer), 250 nM RFC (heteropentamer), 46 μM dATP, 9.7 μM dGTP, 48 μM dCTP, 67 μM dTTP, and either 8.8 nM wild-type pol δ (heterotetramer) or 35 nM exonuclease-deficient pol δ (heterotetramer). Stable assembly (loading) of PCNA onto a P/T junction was not affected by any of the lesions examined in the present or previous study (**Figure S3**)(1). The concentration of each dNTP utilized was within the physiological range observed in dividing human cells (24 ± 22 μM dATP, 5.2 ± 4.5 μM dGTP, 29 ± 19 μM dCTP, 37 ± 30 μM dTTP) (11). Primer extension products were resolved via denaturing polyacrylamide gel electrophoresis, gel images were obtained on a Typhoon Model 9410 imager, and the fluorescence intensity of each DNA band within a given lane was quantified with ImageQuant (GE Healthcare) as described previously (1).

### Primer Extension Analysis

Primer extension assays were analyzed as described previously (1). Briefly, only data points that were less than 20% of the reaction progress (based on the accumulation of primer extension products) were considered. The following analyses apply to all single nucleotide lesions. For analyses of Ab lesions, the corresponding native nucleotide is a guanine (G). The insertion probability is the probability of dNMP incorporation opposite a lesion or the corresponding native nucleotide at dNMP incorporation step *i* and is defined as? P_*i*_. The insertion efficiency is calculated by dividing the insertion probability for a given lesion by the insertion probability for the corresponding native nucleotide in the same sequence context. The extension probability is the probability of dNMP incorporation opposite the 1^st^ template nucleotide 5’ (downstream) of a lesion or the corresponding native nucleotide at dNMP incorporation step *i* and is defined as? P_*i* + 1_. The extension efficiency is calculated by dividing the extension probability for a given lesion by the extension probability for the corresponding native nucleotide in the same sequence context. The bypass probability for a given lesion or the corresponding native nucleotide at dNMP incorporation step *i* is calculated by multiplying the corresponding insertion and extension probabilities (P_*i*_ x P_*i*+1_). The bypass efficiency is calculated by dividing the bypass probability for a given lesion by the bypass probability for the corresponding native nucleotide in the same sequence context. Efficiencies are expressed as percentages which are each obtained by multiplying a given efficiency by 100%.

Encounter of a given lesion or the corresponding native nucleotide at dNMP incorporation step *i* is indicated by the presence of primer extension products resulting from the completion of dNMP incorporation steps up to and including *i*_-1_. These primer extension products are collectively referred to as encounter products. The fraction of progressing pol δ holoenzymes that encounter a given lesion or the corresponding native nucleotide at dNMP incorporation step *i* and subsequently stall (via dissociation of pol δ) at dNMP incorporation step *i* (i.e., during insertion) or *i*+1 (i.e., during extension) is defined as the band intensity of the aborted primer extension product resulting from completion of the prior dNMP incorporation step divided by the sum of the band intensities of all encounter products. These values are utilized to determine the distribution (%) of events that occur after progressing pol δ holoenzymes encounter a lesion or the corresponding native nucleotide at dNMP incorporation step *i*. Specifically, the events are 1) stall during encounter (i.e., stall during insertion); 2) initiate bypass then stall (i.e., complete insertion then stall during extension) and; 3) complete bypass (i.e., complete insertion and extension). The analyses described above were carried out in a corresponding manner for a T<>T lesion or the native di-thymidine sequence that starts at dNMP incorporation *i*. Details of these analyses are provided in the **Supporting Information**.

The contributions (expressed as a percentage) of proofreading by pol δ holoenzymes to insertion opposite a given DNA lesion (i.e., initiation of bypass) was calculated by subtracting the observed insertion efficiency of exonuclease-deficient pol δ holoenzymes from the observed insertion efficiency for wild-type pol δ holoenzymes. The contributions of proofreading by pol δ holoenzymes to (complete) bypass of a given DNA lesion was calculated in an identical manner. Negative values indicate proofreading by pol δ holoenzymes restricts insertion (initiation of bypass) and/or (complete) bypass; positive values indicate that proofreading by pol δ holoenzymes promotes one or both activities.

For all plots in all figures, each data point/column represents the average ± S.E.M. of at least three independent experiments. Error bars are present for all data points on all plots in all figures but may be smaller than the data point. All data presented for 8oxoG, Tg, O6MeG, and εA are from our previous study (1).

## RESULTS

### Lesions resulting from hydrolysis

DNA template nucleobases are susceptible to spontaneous hydrolytic attack at multiple positions. In particular, the N-glycosidic bond connecting the nitrogenous base to the deoxyribose sugar is prone to spontaneous hydrolysis (i.e., depurination or depyrimidination), which generates an Ab lesion. The rate of spontaneous depurination is markedly higher than the rate of spontaneous depyrimidination due to the relative instability of the N-glycosidic bonds of purines. Furthermore, guanines (G) are more likely to undergo spontaneous depurination (**Figure 2A**) than adenines (A) (12). Ab lesions are also generated as intermediates in the base excision DNA repair pathway that repairs small, single-base, non helix distorting lesions resulting from oxidation, deamination, and alkylation (13). Ab lesions are devoid of base-pairing moieties and are the most abundant lesions in human cells, with 10,000 – 20,000 generated per human cell per day (12). Previous biochemical studies indicate that human pol δ predominantly abides by the “A-Rule” during insertion opposite Ab lesions, utilizing dATP for dNMP incorporation (14-17). Thus, within lagging strand DNA templates, abasic sites may be highly mutagenic if progressing pol δ holoenzymes can replicate the lesion. To investigate the effects of G depurination on human lagging strand DNA replication, we characterized and directly compared the progression of pol δ holoenzymes on Cy5-labeled P/T DNA substrates that contain either an abasic site mimic (BioCy5P/T_*i*12_-Ab, **Figure S2**) or a native G (BioCy5P/T_N_, **Figure S2**) 12 nt downstream of the P/T junction (at the 12^th^ dNMP incorporation step, *i*_12_).

**Figure 2.**
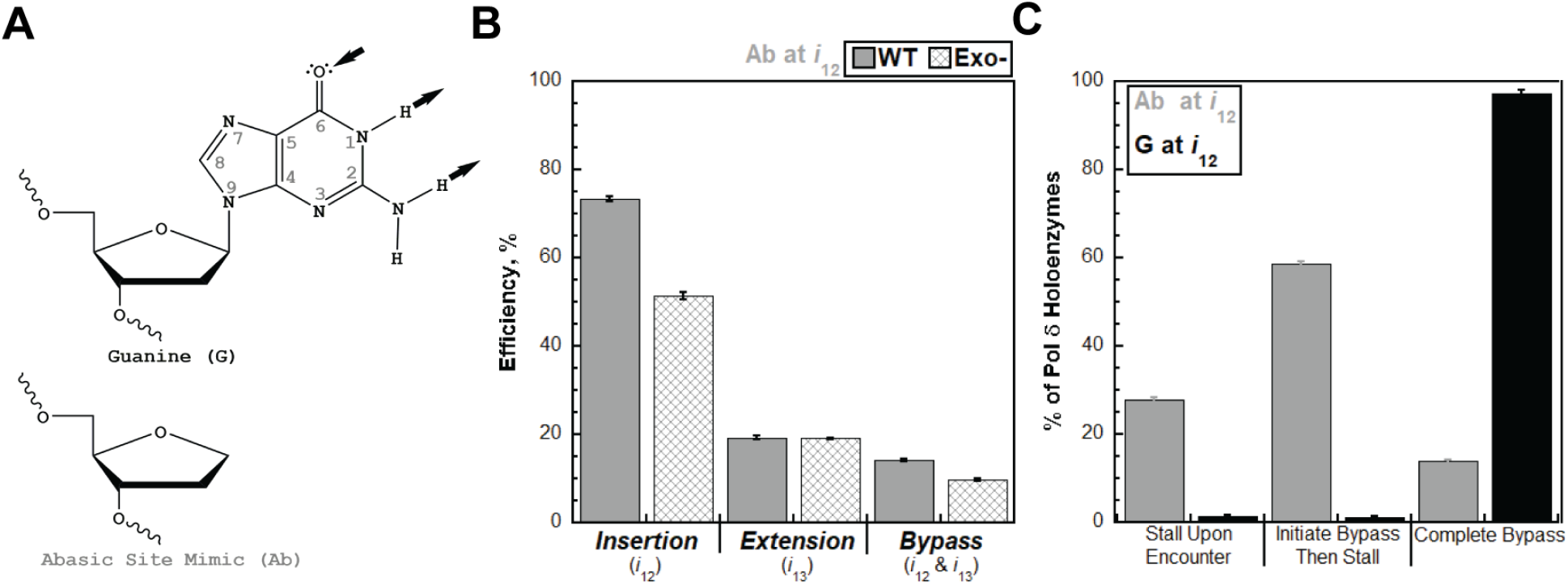
Progressing pol δ holoenzymes encountering an Ab lesion. (**A**) Structure of Ab. Atomic positions of the nitrogenous base are indicated in grey. Canonical base pairing sites within the nucleobase are indicated by black arrows. Arrows pointed towards and away from the nucleobase indicate hydrogen-bonding acceptors and donors, respectively. An Ab is generated from G (*Top*) by cleavage of the N-glycosidic bond connecting position 9 of the nitrogenous base to the deoxyribose sugar. Shown is an Ab mimic (*Bottom*). (**B**) Efficiency of replicating an Ab. The efficiencies of dNMP incorporation opposite an Ab (i.e., insertion), 1 nt downstream of an Ab (i.e., extension), and bypass for wild-type (WT) and exonuclease-deficient (Exo-) pol δ holoenzymes are plotted as percentages. The respective dNMP incorporation step(s) (*i*) for each efficiency is indicated below. (**C**). Fates of pol δ holoenzymes after encountering an Ab. The distribution of pol δ holoenzyme fates observed after an Ab is encountered at *i*_12_ (Ab at *i*_12_) is plotted in Grey. For comparison, the distribution of pol δ holoenzyme fates observed after a native G is encountered at *i*_12_ (G at *i*_12_) is plotted in black.

First, we examined the efficiencies of insertion, extension, and bypass for an Ab lesion (**Figure 2B**). Pol δ holoenzymes are only 14.1 ± 0.4% efficient at bypassing an Ab compared to a native G. The significantly reduced efficiency of Ab bypass is primarily due to a significantly reduced extension efficiency (19.3 ± 0.5%) following a moderately high insertion efficiency (73.3 ± 0.6%). Furthermore, disabling the proofreading activity of pol δ slightly decreases the bypass efficiency (by 4.35 ± 0.42 %) by marginally decreasing the insertion efficiency (by 21.9 ± 1.1 %); the extension efficiency is unaffected. Thus, proofreading during insertion opposite an Ab slightly promotes bypass of this lesion.

Next, we examined the fates of progressing pol δ holoenzymes after encountering an Ab or a native G at *i*_12_ (**Figure 2C**). Nearly all (97.3 ± 0.7 %) pol δ holoenzymes that encounter a native G (G at *i*_12_ in **Figure 2C**) complete bypass prior to stalling, as expected. Only 27.7 ± 0.6 % of pol δ holoenzymes stall upon encountering an Ab (Ab at *i*_12_ in **Figure 2C**) due to the high efficiency of insertion opposite the lesion (*i*_12_, **Figure 2B**). Based on previous biochemical studies, pol δ holoenzymes likely insert dAMP opposite the Ab lesions (14-17). However, more than half (58.5 ± 0.8%) of pol δ holoenzymes that encounter an Ab stall after initiating bypass (**Figure 2C**) due to the significantly reduced efficiency of extension past the lesion (*i*_13_, **Figure 2B**). The remainder (13.8 ± 0.3%) complete bypass of an Ab prior to stalling. Together, these experiments reveal that progressing pol δ holoenzymes contribute to the bypass of a very high proportion (72.3 ± 0.6%) of Ab lesions they encounter, either by initiating bypass (58.5 ± 0.8%) or executing complete bypass (13.8 ± 0.3%). Furthermore, proofreading by pol δ promotes these activities.

The base-pairing moieties at positions 3 and 4 of cytosine (C, **Figure 3A**, *Top*) serve as a hydrogen-bond acceptor and donor, respectively. Position 4 of C is prone to spontaneous hydrolytic attack, replacing the exocyclic primary amine at position 4 with a carbonyl group and changing the nitrogen at position 3 to an amide (18). This process, referred to as hydrolytic deamination, converts cytosine to uracil (U, **Figure 3A**, *Bottom*) and is also catalyzed by cytidine deaminases, such as APOBECs (19). U also forms under nitrosative stress conditions caused by inflammation and after exposure to environmental nitrosative compounds where the exocyclic primary amine at position 4 is attacked directly (i.e., nitrosative deamination)(20). The hydrogen bonding patterns of uracil and thymine (T) are the same where positions 3 and 4 serve as a hydrogen-bond donor and acceptor, respectively. Indeed, eukaryotic DNA polymerases cannot discriminate between a template U and a template T during dNMP incorporation. Thus, while deamination of C only generates 70 – 200 U bases per human cell per day, U is 100% mutagenic if bypassed (18,21). To investigate the effects of C deamination on human lagging strand DNA replication, we characterized and directly compared the progression of pol δ holoenzymes on a Cy5-labeled P/T DNA substrates that contain either a U (BioCy5P/T_*i*11_-U, **Figure S2**) or a native C (BioCy5P/T_N_, **Figure S2**) 11 nt downstream of the P/T junction (at the 11^th^ dNMP incorporation step, *i*_11_).

**Figure 3.**
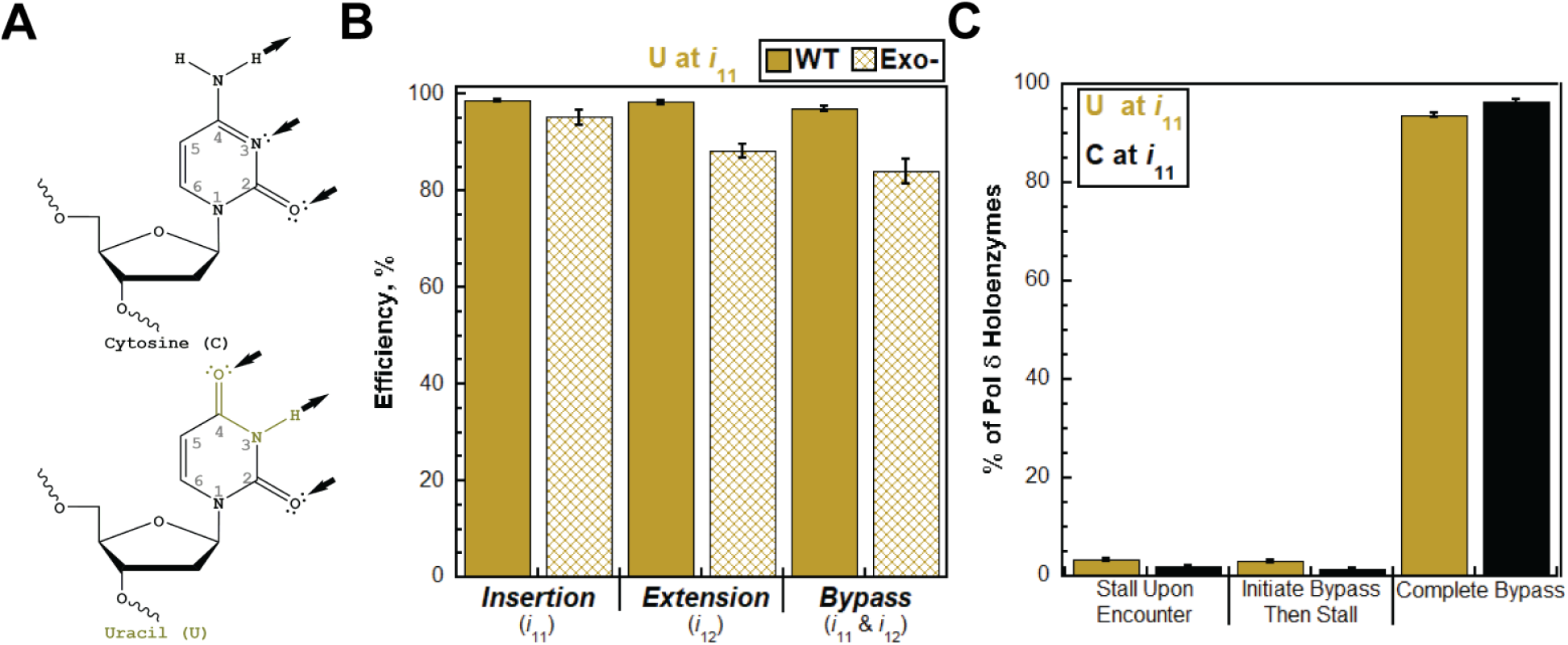
Progressing pol δ holoenzymes encountering a U lesion. (**A**) Structure of U. Atomic positions of the nitrogenous bases are indicated in grey. Canonical base pairing sites within the nucleobase are indicated by black arrows. Arrows pointed towards and away from the nucleobase indicate hydrogen-bonding acceptors and donors, respectively. A U (*Bottom*) is generated from C (*Top*) by deamination of position 4. (**B**) Efficiency of replicating a U. The efficiencies of dNMP incorporation opposite a U (i.e., insertion), 1 nt downstream of a U (i.e., extension), and bypass for wild-type (WT) and exonuclease-deficient (Exo-) pol δ holoenzymes are plotted as percentages. The respective dNMP incorporation step(s) (*i*) for each efficiency is indicated below. (**C**). Fates of pol δ holoenzymes after encountering a U. The distribution of pol δ holoenzyme fates observed after a U is encountered at *i*_11_ (U at *i*_11_) is plotted in Olive. For comparison, the distribution of pol δ holoenzyme fates observed after a native C is encountered at *i*_11_ (C at *i*_11_) is plotted in black.

U does not significantly compromise the efficiencies of insertion (98.7 ± 0.3%) or extension (98.3 ± 0.4%) (**Figure 3B**). Consequently, pol δ holoenzymes are nearly 100% efficient (97.0 ± 0.6%) at bypassing a U lesion compared to a native C (**Figure 3B**). Disabling the proofreading activity of pol δ slightly decreases the bypass efficiency (by 13.0 ± 2.5 %), primarily by slightly decreasing the extension efficiency (by 10.0 ± 1.4 %). The insertion efficiency is minimally decreased (by 3.5 ± 1.6%). Thus, proofreading promotes bypass of U by promoting extension past the lesion. This may occur by proofreading insertion of an incorrect dNMP opposite U (mismatch) to promote extension and/or proofreading extension to stabilize correct dNMP incorporation 1 nt downstream of U. These alternatives cannot be discerned under the conditions of the assay.

Only 6.34 ± 0.44 % of pol δ holoenzymes that encounter a U at *i*_11_ stall prior to bypassing the lesion (**Figure 3C**) due to significantly high efficiencies of insertion and extension for the lesion (**Figure 3B**). Specifically, 3.30 ± 0.23 % stall upon encounter and 3.04 ± 0.25 % initiate bypass and then stall. The distribution of these stalling events is nearly identical to that observed for pol δ holoenzymes that encounter a native C at *i*_11_ (**Figure 3C**). Thus, stalling of pol δ holoenzymes is not significantly promoted by encounters with U lesions relative to native C. These experiments reveal that the progression of pol δ holoenzymes is not significantly affected by encounters with U lesions and, consequently, 93.7 ± 0.4 % of U lesions encountered by pol δ holoenzymes are bypassed prior to stalling. Proofreading by pol δ promotes this activity. Together with the miscoding potential of U described above, this suggests that U lesions in lagging strand templates are highly mutagenic, generating C→T transition mutations.

The base-pairing moieties at positions 1 and 6 of adenine (A, **Figure 4A**, *Top*) serve as a hydrogen-bonding acceptor and donor, respectively. The N6-amine of A is also prone to hydrolytic and nitrosative deamination, which converts A to inosine (I, **Figure 4A**, *Bottom*) (18). (22). Inosine presents a hydrogen-bonding pattern at N1 and C6 that is identical to that of G. Indeed, biochemical studies indicate that I is primarily interpreted by DNA polymerases as a template G (23,24). Thus, I lesions within lagging strand DNA templates may be highly mutagenic, promoting A→G transition mutations if replicated by progressing pol δ holoenzymes. To investigate the effects of A deamination on human lagging strand DNA replication, we characterized and directly compared the progression of pol δ holoenzymes on Cy5-labeled P/T DNA substrates that contain either an I (BioCy5P/T_*i*10_-I, **Figure S2**) or a native A (BioCy5P/T_N_, **Figure S2**) 10 nt downstream of the P/T junction (at the 10^th^ dNMP incorporation step, *i*_10_).

**Figure 4.**
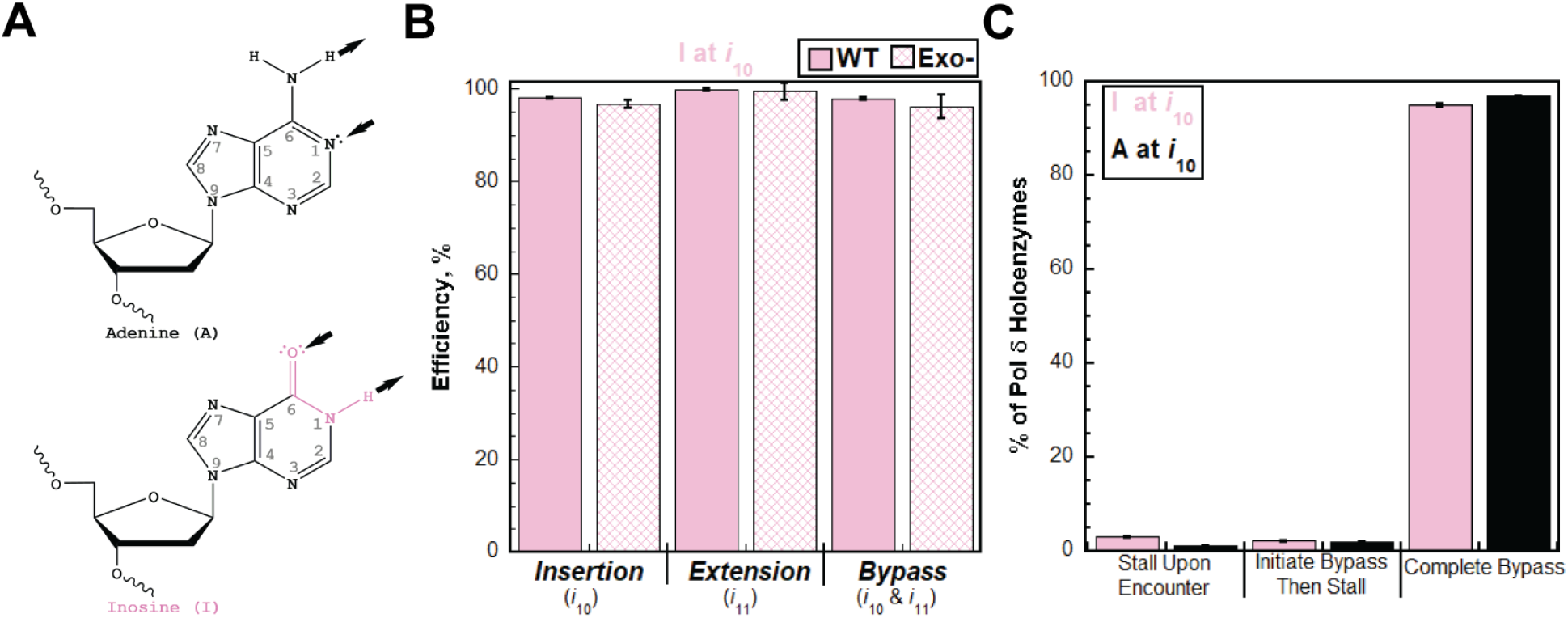
Progressing pol δ holoenzymes encountering an I lesion. (**A**) Structure of I. Atomic positions of the nitrogenous bases are indicated in grey. Canonical base pairing sites within the nucleobase are indicated by black arrows. Arrows pointed towards and away from the nucleobase indicate hydrogen-bonding acceptors and donors, respectively. An I (*Bottom*) is generated from an A (*Top*) by deamination of position 6. (**B**) Efficiency of replicating an I. The efficiencies of dNMP incorporation opposite a U (i.e., insertion), 1 nt downstream of a U (i.e., extension), and bypass for wild-type (WT) and exonuclease-deficient (Exo-) pol δ holoenzymes are plotted as percentages. The respective dNMP incorporation step(s) (*i*) for each efficiency is indicated below. (**C**). Fates of pol δ holoenzymes after encountering an I. The distribution of pol δ holoenzyme fates observed after an I is encountered at *i*_10_ (I at *i*_10_) is plotted in Pink. For comparison, the distribution of pol δ holoenzyme fates observed after a native A is encountered at *i*_10_ (A at *i*_10_) is plotted in black.

The insertion (98.0 ± 0.2%) and extension efficiencies (99.8 ± 0.2%) of pol δ holoenzymes are insignificantly compromised by I, if at all (**Figure 4B**). Consequently, pol δ holoenzymes are nearly 100% efficient (97.8 ± 0.4%) at bypassing an I lesion compared to a native A (**Figure 4B**). Disabling the proofreading activity of pol δ had no effect on the efficiencies of insertion or extension, indicating that this activity does not contribute to the bypass of I. Finally, only 5.10 ± 0.24 % of pol δ holoenzymes that encounter I at *i*_10_ stall prior to bypassing the lesion (**Figure 4C**) due to the near-maximum efficiencies of insertion and extension for the lesion (**Figure 4B**). Specifically, 2.96 ± 0.17 % upon encounter and 2.14 ± 0.16 % initiate bypass and then stall (**Figure 4C**). The distribution of these stalling events is essentially identical to that observed for pol δ holoenzymes that encounter a native A at *i*_10_. Thus, I lesions do not promote stalling of pol δ holoenzymes relative to native A. As was observed for U, these experiments reveal that the progression of pol δ holoenzymes is essentially unaffected by encounters with I lesions and, consequently, 94.9 ± 0.2 % of I lesions are bypassed prior to stalling. Considering the miscoding potential of I described above, this suggests that I lesions in lagging strand templates are highly mutagenic, generating A→G transition mutations.

### A Lesion resulting from oxidation

Guanine (G, **Figure 5A**, *Top*), because of its relatively low oxidation potential, is the template DNA nucleobase that is most susceptible to oxidation from reactive oxygen species (ROS) (25-28). Specifically, position 8 of G is susceptible to attack by a hydroxyl radical and β-fragmentation of the imidazole ring within the resultant unstable intermediate, followed by reduction, yields *N*6-(2-deoxy-α,β-D-erythro-pentofuranosyl)-2,6-diamino-4-hydroxy-5-formamido-pyrimidine (FapyG, **Figure 5A**, *Bottom*)(29,30). FapyG has an unusual ability to isomerize (via epimerization) between α- and β-anomers under physiological conditions. A recent study suggests that FapyG in template ssDNA exists in a ∼1:1 anomeric ratio (29).

**Figure 5.**
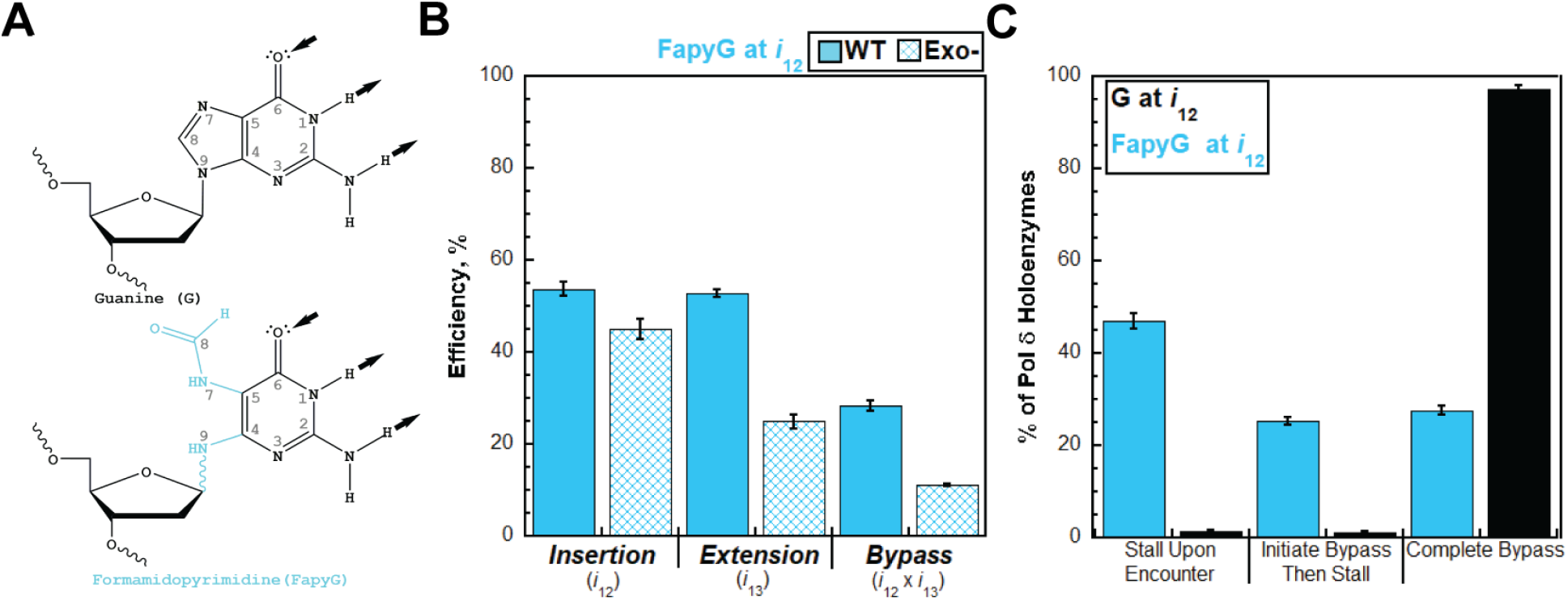
Progressing pol δ holoenzymes encountering a FapyG lesion. (**A**) Structure of FapyG. Atomic positions of the nitrogenous bases are indicated in grey. Canonical base pairing sites within the nucleobase are indicated by black arrows. Arrows pointed towards and away from the nucleobase indicate hydrogen-bonding acceptors and donors, respectively. A FapyG (*Bottom*) is generated from an G (*Top*) by β-fragmentation of the imidazole ring at position 6. The N-glycosidic bond of FapyG is displayed as a wave to highlight the anomers of this lesion where the bond is either in the α- or β-position. (**B**) Efficiency of replicating a FapyG. The efficiencies of dNMP incorporation opposite a FapyG (i.e., insertion), 1 nt downstream of a FapyG (i.e., extension), and bypass for wild-type (WT) and exonuclease-deficient (Exo-) pol δ holoenzymes are plotted as percentages. The respective dNMP incorporation step(s) (*i*) for each efficiency is indicated below. (**C**). Fates of pol δ holoenzymes after encountering a FapyG. The distribution of pol δ holoenzyme fates observed after a FapyG is encountered at *i*_12_ (FapyG at *i*_10_) is plotted in Cyan. For comparison, the distribution of pol δ holoenzyme fates observed after a native G is encountered at *i*_12_ (G at *i*_12_) is plotted in black.

Depending on the level and type of oxidative stress, the cellular levels of FapyG are comparable to or even greater than 8-oxo-7,8-dihydro-2′-deoxyguanosine (8OxoG), another common lesion derived from G oxidation at position 8 (31-35). Like 8OxoG, the properties of the base-pairing moieties at positions 1, 2, and 6 of FapyG (i.e., the Watson-Crick face) are identical to those in native G (**Figure 5A**). FapyG and 8OxoG lesions are replicated by DNA polymerases in a predominantly error-free manner but both promote erroneous insertion of dAMP opposite these lesions, leading to G→T transversion mutations (14,25,36-39).

Interestingly, FapyG is more mutagenic than 8OxoG in mammalian cells (36,37) and the fidelities of replicating each lesion are governed by distinct interactions within the polymerase active site. DNA polymerases engage 8OxoG lesions in either the *anti* or *syn* conformation. In the *anti*-conformation, the Watson-Crick face of 8OxoG is presented to base pair with an incoming dCTP. In the *syn*-conformation, the Hoogsteen face of 8OxoG is presented to base pair with an incoming dATP (14,25,39). For FapyG, both anomers are engaged by DNA polymerases in the *anti*-conformation where the Watson-Crick face of the lesion is presented to base pair with an incoming dNTP. The β-anomer promotes efficient insertion of dCMP. The α-anomer promotes inefficient insertion of either dCMP or dAMP (29). These precedents suggest that FapyG and 8OxoG lesions within lagging strand DNA templates promote G→T transversion mutations if replicated by progressing pol δ holoenzymes. The effects of 8oxoG on human lagging strand replication were analyzed in our previous study (1). To investigate the effects of FapyG lesions on human lagging strand DNA replication, we characterized and directly compared the progression of pol δ holoenzymes on Cy5-labeled P/T DNA substrates that contain either a FapyG (BioCy5P/T_*i*12_-FapyG, **Figure S2**) or a native G (BioCy5P/T_N_, **Figure S2**) 12 nt downstream of the P/T junction (at the 12^th^ dNMP incorporation step, *i*_12_).

The insertion (53.8 ± 1.7%) and extension efficiencies (52.8 ± 0.8 %) of pol δ holoenzymes are both reduced ∼50% by FapyG, (**Figure 5B**). Consequently, pol δ holoenzymes are only 28.4 ± 1.1 % efficient at bypassing a FapyG compared to a native G (**Figure 5B**). The ∼50% reductions in the insertion and extension efficiencies could reflect the anomeric ratio of FapyG in the template ssDNA, where the α-anomer accounts for ∼50% of the lesions and is inefficiently bypassed by DNA polymerases (29). Furthermore, disabling the proofreading activity of pol δ significantly decreases the bypass efficiency (by 17.2 ± 1.13 %) primarily by decreasing the extension efficiency (by 27.9 ± 1.7 %), as the insertion efficiency is decreased by a smaller amount (8.78 ± 2.73%). Thus, the 3′→5′ exonuclease activity of human pol δ promotes FapyG bypass by proofreading insertion opposite the lesion and promoting extension beyond the lesion. Again, the latter may be achieved by proofreading insertion of an incorrect dNMP opposite FapyG (mismatch) to promote extension and/or proofreading extension to stabilize correct dNMP incorporation 1 nt downstream of FapyG. These alternatives cannot be discerned under the conditions of the assay.

Less than half (47.0 ± 1.6 %) of pol δ holoenzymes stall upon encountering FapyG (FapyG at *i*_12_ in **Figure 5C**) due to the moderate efficiency of insertion opposite the lesion (*i*_12_, **Figure 5B**). Based on previous studies of DNA polymerases, pol δ holoenzymes preferentially insert dCMP (over dAMP) opposite the lesion (29,38). However, due to the significantly reduced efficiency of extension past the lesion (*i*_13_, **Figure 5B**), approximately one-fourth of pol δ holoenzymes (25.3 ± 0.7 %) that encounter a FapyG initiate bypass and then stall (**Figure 5C**). The remainder (27.6 ± 1.0 %) complete bypass prior to stalling. Together, these experiments reveal that progressing pol δ holoenzymes contribute to the bypass of more than half (53.0 ± 1.6%) of the FapyG lesions they encounter; either by initiating bypass (25.3 ± 0.7 %) or executing complete bypass (27.6 ± 1.0 %). Proofreading by pol δ promotes these activities. Interestingly, the contributions of pol δ holoenzymes to the bypass of FapyG are distinct from those for 8OxoG. Our previous study revealed that progressing pol δ holoenzymes contribute to the bypass of a very high proportion (84.2 ± 0.6%) of the 8OxoG lesions they encounter; nearly half (47.8 ± 0.8%) are replicated by pol δ holoenzymes (i.e., insertion only) and more than a one-third (36.4 ± 1.5 %) are completely bypassed (1). The results for 8OxoG are displayed and discussed further in the **Discussion** below. These direct comparisons reveal that FapyG lesions are more significant impediments to pol δ holoenzyme progression than 8OxoG lesions in the same sequence context.

### A Lesion resulting from alkylation

The nitrogen group at position 3 of C (**Figure 6A**, *Top*) serves as a hydrogen-bond acceptor and is susceptible to attack by S_N_2 alkylating agents when C is exposed in ssDNA during DNA metabolic processes, such as transcription. In the resultant lesion, 3-methyl-2’-deoxycytidine (3MeC, **Figure 6A**, *Bottom*), a methyl group is covalently attached to the nitrogen at position 3, removing its ability to hydrogen bond and introducing a positive charge on the nitrogenous base (40,41). 3MeC lesions are unable to base pair and, hence, have been viewed as strong blocks to DNA replication. Indeed, only ∼15 – 20% of 3MeC lesions in ssDNA vectors are bypassed by replicative DNA polymerases in *E coli* that are deficient in DNA repair and the DNA damage response. Furthermore, bypass of 3MeC in this context primarily resulted in C→A transversion and C→T transition mutations, indicating that *E. coli* replicative DNA polymerases utilize either dTTP or dATP for insertion opposite 3MeC (42). Finally, a recent *in vitro* study revealed that excess human pol δ in isolation catalyzes insertion of primarily dTMP and dAMP opposite 3MeC over long incubation times but bypass is very limited (40). Altogether, this suggests that 3MeC lesions in lagging strand DNA templates may promote C→A transversion and C→T transition mutations if replicated by progressing pol δ holoenzymes. To investigate the effects of 3MeC lesions on human lagging strand DNA replication, we characterized and directly compared the progression of pol δ holoenzymes on Cy5-labeled P/T DNA substrates that contain either a 3MeC (BioCy5P/T_*i*11_-3MeC, **Figure S2**) or a native C (BioCy5P/T_N_, **Figure S2**) 11 nt downstream of the P/T junction (at the 11^th^ dNMP incorporation step, *i*_11_).

**Figure 6.**
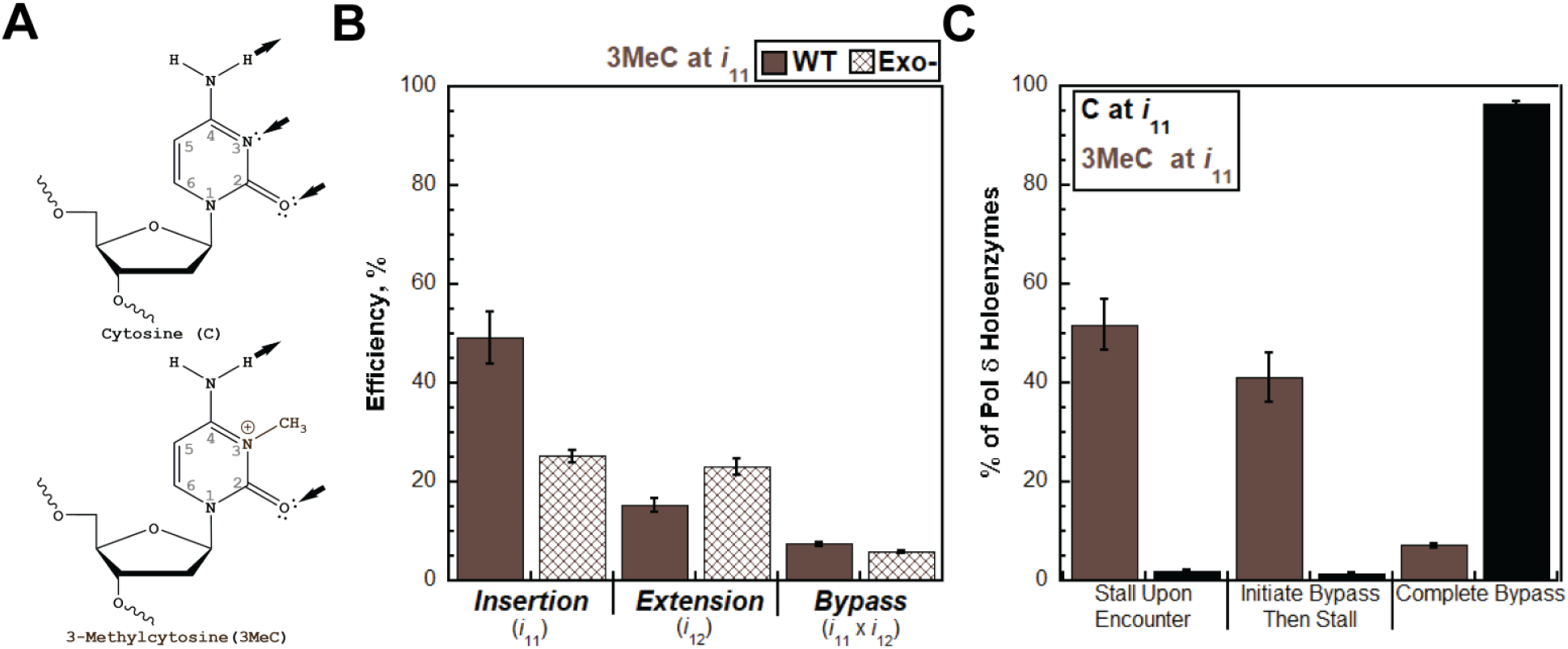
Progressing pol δ holoenzymes encountering a 3MeC lesion. (**A**) Structure of 3MeC. Atomic positions of the nitrogenous bases are indicated in grey. Canonical base pairing sites within the nucleobase are indicated by black arrows. Arrows pointed towards and away from the nucleobase indicate hydrogen-bonding acceptors and donors, respectively. A 3MeC (*Bottom*) is generated from an C (*Top*) by methylation of position 3. (**B**) Efficiency of replicating a 3MeC. The efficiencies of dNMP incorporation opposite a 3MeC (i.e., insertion), 1 nt downstream of a 3MeC (i.e., extension), and bypass for wild-type (WT) and exonuclease-deficient (Exo-) pol δ holoenzymes are plotted as percentages. The respective dNMP incorporation step(s) (*i*) for each efficiency is indicated below. (**C**). Fates of pol δ holoenzymes after encountering a 3MeC. The distribution of pol δ holoenzyme fates observed after a 3MeC is encountered at *i*_11_ (3MeC at *i*_11_) is plotted in Brown. For comparison, the distribution of pol δ pol δ holoenzyme fates observed after a native C is encountered at *i*_11_ (C at *i*_11_) is plotted in black.

Pol δ holoenzymes are only 7.39 ± 0.46 % efficient at bypassing a 3MeC compared to a native C (**Figure 6B**). The significantly reduced efficiency of 3MeC bypass is primarily due to a significantly reduced extension efficiency (15.3 ± 1.5%) following a moderate insertion efficiency (49.2 ± 5.3%). Interestingly, disabling the proofreading activity of pol δ significantly decreases the insertion efficiency (by 24.0 ± 5.5 %) but slightly increases the extension efficiency (by 7.88 ± 2.22 %). This results in a slight reduction in the bypass efficiency (by 1.59

± 0.52%). Thus, proofreading contributes only marginally to the overall bypass of 3MeC lesions due to counteracting effects of pol δ’s 3’→5’ exonuclease activity; proofreading promotes insertion opposite 3MeC but restricts extension past this lesion. These observations are unique among the lesions analyzed in the previous and present studies and suggest that proofreading by pol δ restricts holoenzyme progression after initiating 3MeC bypass.

Approximately half (51.8 ± 5.2%) of pol δ holoenzymes stall upon encountering 3MeC (3MeC at *i*_11_ in **Figure 6C**) due to the moderate efficiency of insertion opposite the lesion (*i*_11_, **Figure 6B**). Based on a previous study, pol δ holoenzymes likely insert dTMP or dAMP opposite the lesion (40). However, due to the significantly reduced efficiency of extension past this lesion (*i*_12_, **Figure 6B)**, 41.1 ± 5.0 % of pol δ holoenzymes that encounter 3MeC initiate bypass and then stall (**Figure 6C**). Only 7.13 ± 0.45% complete bypass prior to stalling.

Together, these experiments reveal that progressing pol δ holoenzymes contribute to the bypass of approximately half (48.2 ± 5.2 %) of the 3MeC lesions they encounter, either by initiating bypass (41.5 ± 5.0 %) or executing complete bypass (7.13 ± 0.45%). Furthermore, proofreading by pol δ promotes initiation of bypass but restricts further holoenzyme progression.

### A Lesion resulting from ultraviolet radiation-induced intrastrand crosslinking

Template DNA nucleobases have absorption maximum in the ultraviolet (UV) wavelength region and, hence, directly absorb solar ultraviolet radiation (UVR), UVR-B (280 – 315 nm) in particular. A consequence is the generation of lesions in the form of pyrimidine photoproducts. Cyclobutane pyrimidine dimers (CPDs) constitute ∼75% of the UVR-induced lesions and are generated via intrastrand crosslinking of positions 5 and 6 of adjacent pyrimidine nucleobases where one pyrimidine nucleobase is in the *syn-*conformation and the other is the *anti*-conformation (43,44).

Shown in **Figure 7A** is a CPD (T<>T, *right* panel) generated from adjacent thymine (T) nucleobases (*left* panel). CPDs, such as T<>T, are viewed as complete blocks to the progression of human canonical DNA replication pathways, which employ the B-family, high-fidelity DNA polymerases ε and δ for leading and lagging strand DNA replication, respectively (5). However, due to the scarcity of studies, it is unknown where canonical DNA replication pathways stall in relation to CPDs (16,45). To investigate the effects of CPDs on human lagging strand DNA replication, we characterized and directly compared the progression of pol δ holoenzymes on Cy5-labeled P/T DNA substrates that contain either a T<>T (BioCy5P/T_*i*11-12_-T<>T, **Figure S2**) or a native TT sequence (BioCy5P/T_N*_, **Figure S2**) 11 nt downstream of the P/T junction (at the 11^th^ and 12^th^ dNMP incorporation steps, *i*_11_ and *i*_12_). For this lesion, bypass requires insertion opposite the 3’ T within the T<>T (i.e., insertion 1), insertion opposite the 5’ T (i.e., insertion 2), and extension past the 5’ T.

**Figure 7.**
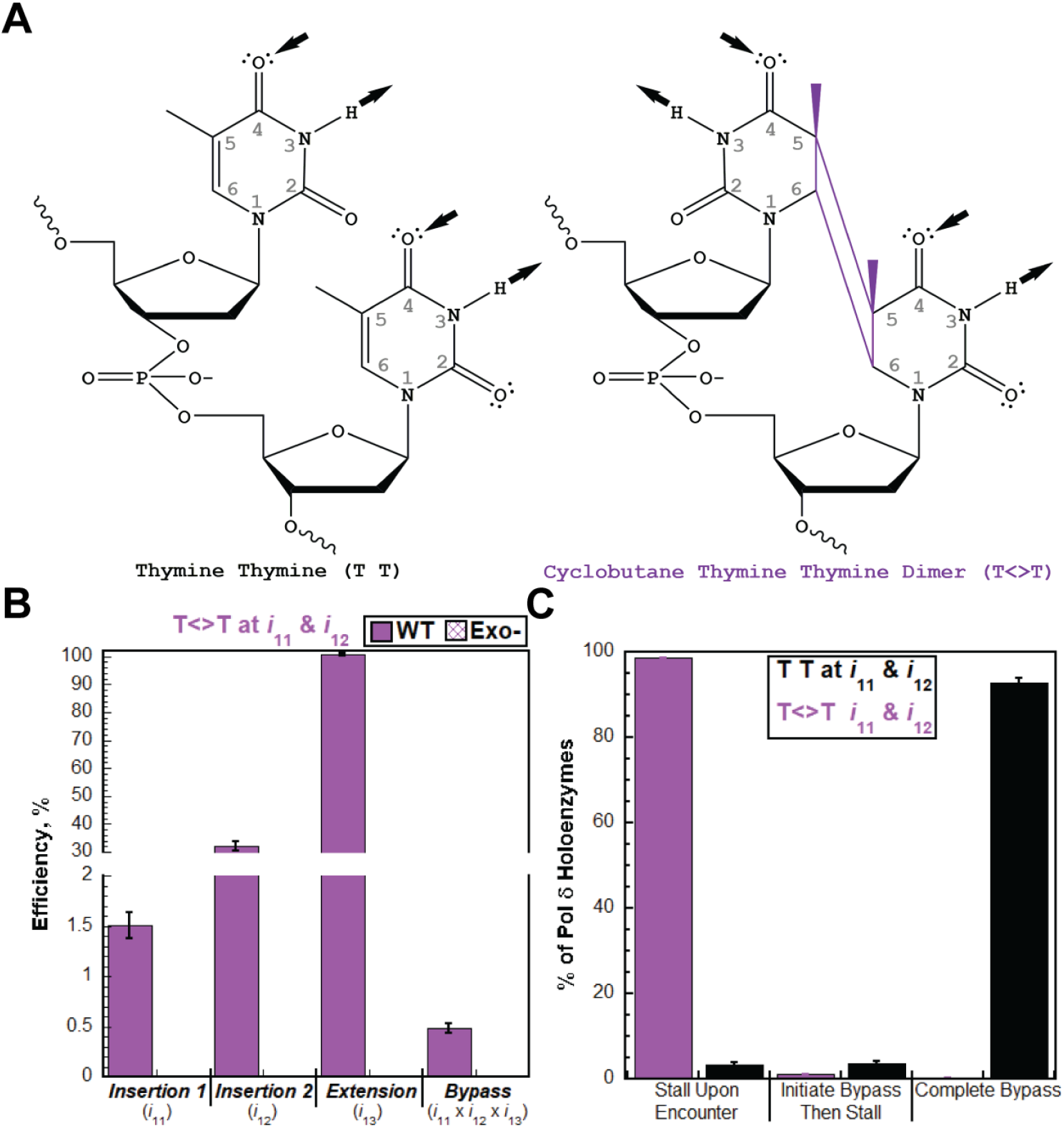
Progressing pol δ holoenzymes encountering a T<> lesion. (**A**) Structure of T<>T. Atomic positions of the nitrogenous bases are indicated in grey. Canonical base pairing sites within the nucleobases are indicated by black arrows. Arrows pointed towards and away from the nucleobase indicate hydrogen-bonding acceptors and donors, respectively. A T<>T (*Right*) is generated from an T T sequence (*Left*) by crosslinking the 5 positions and the 6 positions of adjacent thymines via covalent carbon-carbon bonds. (**B**) Efficiency of replicating a T<>T. The efficiencies of dNMP incorporation opposite the 5’ nucleotide in a T<>T (i.e., insertion 1), opposite the 3’ nucleotide in a T<>T (i.e., insertion 2), 1 nt downstream of a T<>T (i.e., extension), and bypass for wild-type (WT) and exonuclease-deficient (Exo-) pol δ holoenzymes are plotted as percentages. Insertion 1, insertion 2, extension, and bypass were not observed for exopol δ holoenzymes. The respective dNMP incorporation step(s) (*i*) for each efficiency is indicated below. (**C**). Fates of progressing pol δ holoenzymes after encountering a T<>T. The distribution of pol δ holoenzyme fates observed after a T<>T is encountered at *i*_11_ and *i*_12_ (T<>T at *i*_11 &_ *i*_12_) is plotted in purple. For comparison, the distribution of pol δ holoenzyme fates observed after a native T T sequence is encountered at *i*_11_ and *i*_12_ (T T at *i*_11 &_ *i*_12_) is plotted in black.

Pol δ holoenzymes are only 0.483 ± 0.048 % efficient at bypassing a T<>T compared to a native TT sequence (**Figure 7B**). The near-zero bypass efficiency is essentially due to an almost complete elimination of the insertion 1 efficiency (1.51 ± 0.13 %). Interestingly, the efficiencies of insertion 2 (32.1 ± 1.9 %) and extension past the T<>T (100 ± 0.502 %) are each significant. However, less than 1.45 ± 0.13 % and 0.457 ± 0.048 % of progressing pol δ holoenzymes that encounter a T<>T reach the insertion 2 and extension steps (**Figure 7C**), respectively. This renders the surprisingly high efficiencies for these dNMP incorporation steps futile. Disabling the proofreading activity of human pol δ eliminates the minor dNMP incorporation observed for insertion 1 (1.45 ± 0.13 %) and consequently all subsequent dNMP incorporation events downstream. Thus, proofreading by pol δ does not deter bypass of the lesion. Finally, nearly all pol δ holoenzymes (98.5 ± 0.1%) stall upon encountering a T<>T (“T<>T at *i*_11 &_ *i*_12_” in **Figure 7C**) due to the near-zero efficiency of insertion 1 (*i*_11_, **Figure 7B**). A minor percentage of pol δ holoenzymes (1.08 ± 0.13%) that encounter a T<>T initiate bypass and then stall, either during insertion 2 (1.00 ± 0.09 %) or extension (0.082 ± 0.037 %). Only 0.37 ± 0.03 % complete bypass prior to stalling. Altogether, the results from these analyses reveal that essentially all pol δ holoenzymes stall upon encountering a T<>T and this is not promoted by pol δ proofreading.

## Discussion

We recently reconstituted human lagging strand replication using a fluorescence primer extension assay to quantitatively analyze, at single nucleotide resolution, the enzymatic activities of pol δ holoenzymes during initial binding events with P/T DNA substrates that each contain a prominent lesion resulting from alkylation or oxidation (1). P/T DNA substrates contained either a single 8OxoG, Tg, O6MeG, or εA lesion ≥ 9 nt downstream of the P/T junction from which DNA synthesis initiates. This assay directly reports on; 1) the efficiencies of the dNMP incorporation steps comprising lesion bypass by pol δ holoenzymes; 2) the distribution of events observed after progressing pol δ holoenzymes encounter a lesion and; 3) the contribution of proofreading by pol δ to lesion bypass. This initial study provided seminal insights into the replication of damaged lagging strand templates, challenging our fundamental understanding of lagging strand DNA replication and DDT. In the present study, we expand the scope of this investigation to other biologically significant, structurally diverse lesions. Collectively, examination of ten lesions from the previous and present study (**Figure 8A**) reveal that the efficiencies of insertion and bypass for human pol δ holoenzymes each span a complete range (from 0 to 100 %, **Figure 8B**). Consequently, the fates of progressing pol δ holoenzymes upon encountering lesions are quite diverse (**Figure 8C**). Furthermore, pol δ proofreading does not deter bypass of any lesion analyzed. Rather, it significantly promoted bypass for 7 of the 10 lesions analyzed (**Figure 9**). These results together with those from previous reports on human pol δ, significantly alter our understanding of the replicative capacity of high-fidelity pol δ?holoenzymes and their functional role(s) in lesion bypass on lagging strand templates, as discussed further below.

**Figure 8.**
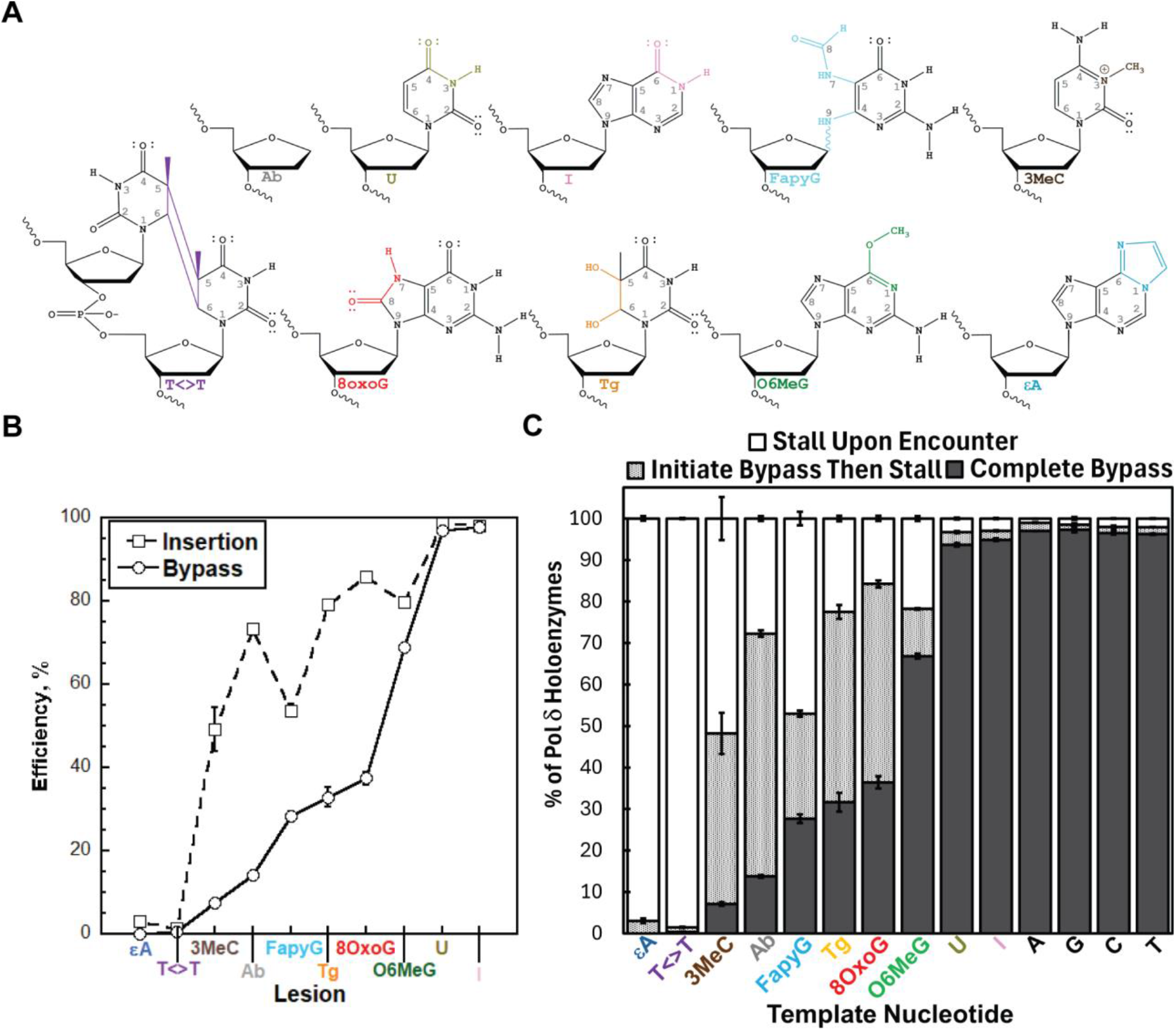
Progressing pol δ holoenzymes encountering DNA lesions. (**A**) Structures of the DNA lesions analyzed in our previous and present studies. For each, the covalent modifications are uniquely color-coded. (**B**) Efficiency of replicating and bypassing DNA lesions. The efficiencies of insertion and bypass for DNA lesions analyzed in our previous and present studies are plotted. For T<>T, the efficiency of dNMP incorporation opposite the 5’ nucleotide within the CPD (i.e., insertion 1) is plotted. (**C**). Distributions of pol δ holoenzyme fates observed after damaged or native template nucleotides are encountered. For each, the % distribution of pol δ holoenzymes that stall upon encounter, initiate bypass then stall, and complete bypass is displayed as a stacked column where the sum of the percentages for all events is equal to 100%. The data for Ab, U, I, FapyG, 3MeC, and T<>T lesions and G, C, and A native template nucleotides are from **Figures 2C, 3C, 4C, 5C, 6C**, and **7C** from the main text. The data for εA, Tg, O6MeG, and 8OxoG lesions and a native T template nucleotide are from our previous study (1).

**Figure 9.**
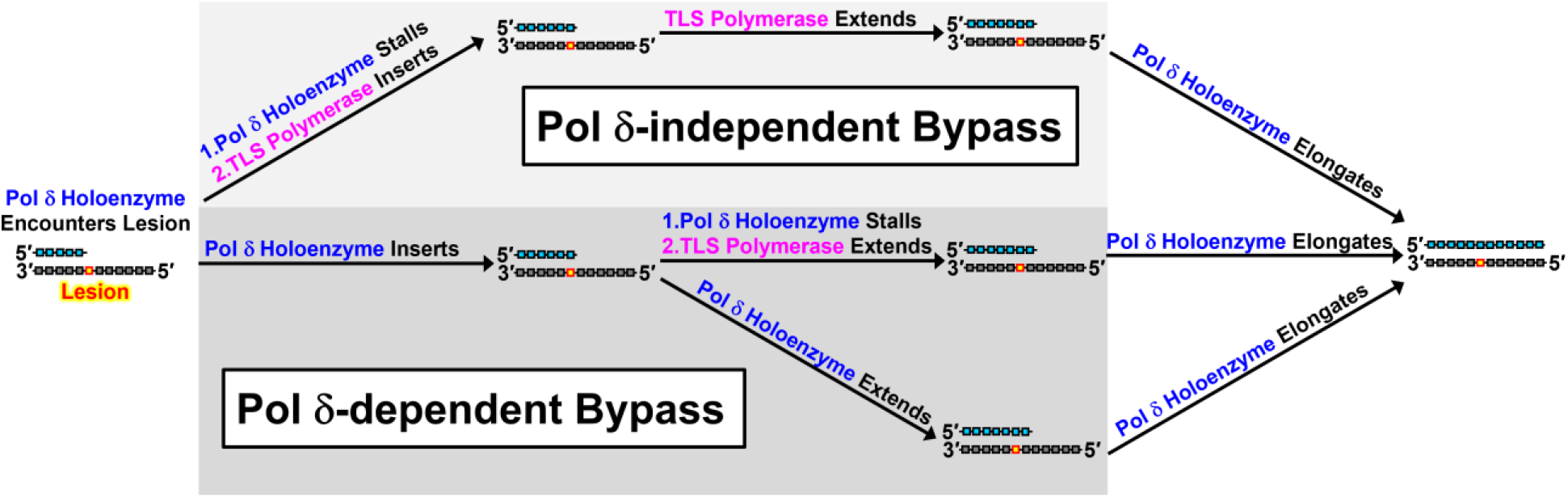
3MeC, FapyG, Ab, Tg, and 8OxoG lesions in lagging strand templates are bypassed by pol δ-independent and -dependent mechanisms. Only P/T DNA is shown for simplicity. Nascent (primer) and template DNA strands are depicted in cyan and grey, respectively. *(Top)* Pol δ-independent lesion bypass. (*Bottom*) Pol δ-dependent lesion bypass mechanisms.

**Figure 9.**
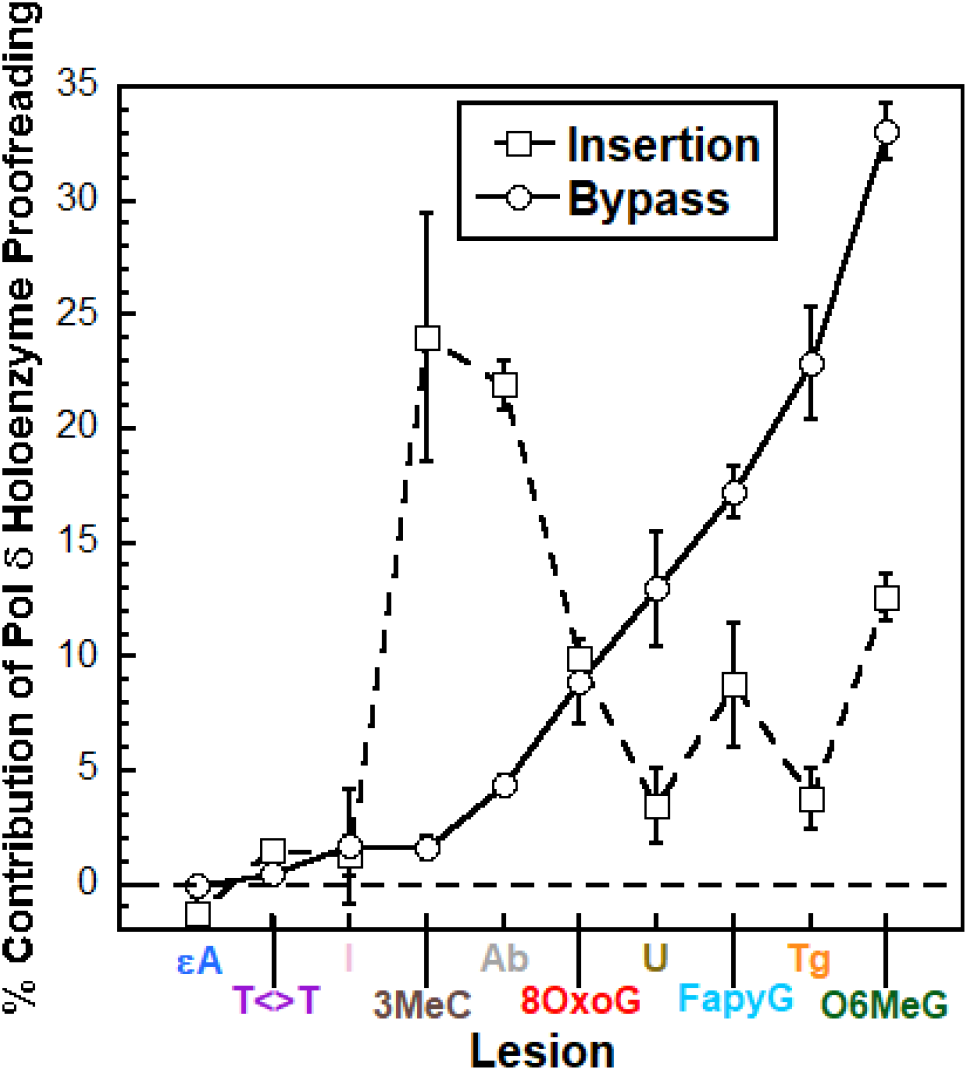
Proofreading by pol δ holoenzymes and lesion bypass. The % contribution of proofreading by pol δ holoenzymes to insertion opposite lesions (i.e., initiation of bypass) and (complete) bypass is plotted. The data for Ab, U, I, FapyG, 3MeC, and T<>T lesions are calculated from **Figures 2B, 3B, 4B, 5B, 6B**, and **7B** from the main text. The data for εA, Tg, O6MeG, and 8OxoG lesions are from our previous study (1). Negative values, which are not observed, indicate that proofreading by pol δ holoenzymes restricts insertion and/or bypass; positive values indicate that proofreading by pol δ holoenzymes promotes insertion and/or bypass.

### In lagging strand templates, a high proportion of O6MeG lesions and nearly all U and I lesions are bypassed by pol δ holoenzymes

Human pol δ holoenzymes are remarkably efficient at insertion opposite and extension beyond O6MeG lesions (**Figure 8B**). Consequently, 66.8 ± 0.6% of progressing pol δ holoenzymes that encounter O6MeG lesions execute complete bypass prior to stalling (**Figure 8C**). This indicates that progressing pol δ holoenzymes bypass more than two-thirds of the O6MeG lesions they encounter. Based on these results, we propose that a high proportion of O6MeG lesions in lagging strand templates are bypassed by progressing pol δ holoenzymes, rather than a DDT pathway. This agrees with results from numerous independent studies of human cells that collectively suggest that O6MeG lesions are bypassed by pol δ holoenzymes (67-70).

Human pol δ holoenzymes are nearly 100% efficient at insertion opposite and extension beyond U and I lesions (**Figure 8B**). Consequently, ≥ 93.7 ± 0.4 % of progressing pol δ holoenzymes that encounter these lesions execute complete bypass prior to stalling (**Figure 8C**). These values are nearly identical to those observed for bypass of the corresponding native template nucleotides (C for U, A for I) in the same sequence context (**Figure 8C**). This indicates that progressing pol δ holoenzymes do not recognize U or I as lesions and, consequently, bypass nearly all U and I template nucleotides they encounter. Based on these results, we propose that all U and I in lagging strand templates are bypassed by progressing pol δ holoenzymes, rather than a DDT pathway. Accordingly, U and I that persist in lagging strand templates during S-phase are significant mutagenic threats to genome integrity.

The results from the present study on the bypass of U by human pol δ holoenzymes (**Figure 8B – C**) agree with a recent report that demonstrated that both pol δ and pol ε holoenzymes from *S. Cerevisae* also do not recognize U as a lesion and readily bypass it (21). Pol δ and pol ε are responsible for the replication of lagging and leading strand templates, respectively (71-75). Interestingly, *in vitro* and *in vivo* studies from a recent report suggest that U lesions stall progression of human pol ε holoenzymes and, hence, leading strand replication, ultimately activating DDT pathways (76). This suggests that human pol ε holoenzymes recognize U as a lesion, in contrast to pol ε holoenzymes from *S. cerevisiae*. Together with the findings from the present study, this also suggests that U lesions in the human genome are a selective impediment to leading strand replication, as they are readily bypassed by progressing pol δ holoenzymes (**Figure 8B - C**). To the best of our knowledge, studies on the bypass of I lesions in human cells or by pol δ holoenzymes from any organism have yet to be reported.

### εA and T<>T are strong blocks to pol δ holoenzyme progression

Previous *ex vivo* and *in vivo* studies of human cells revealed that DDT pathways are required for the bypass of essentially all εA and T<>T lesions (**Figure 8A**) in lagging strand templates (46-52). However, due to the scarcity of studies, it is unknown where pol δ holoenzymes stall in relation to these lesions and what functional roles, if any, pol δ holoenzymes serve in the bypass of these lesions. Our previous and present study reveal that human pol δ holoenzymes are drastically inefficient at insertion opposite εA and the 3’ T within a T<>T (**Figure 8B**)(1).

Consequently, nearly all progressing pol δ holoenzymes (≥ 96.9 ± 0.6 %) stall upon encountering an εA or a T<>T lesion in lagging strand templates (**Figure 8C**). The minor percentage of progressing pol δ holoenzymes that initiate bypass of these lesions via insertion (≤ 3.11 ± 0.57 %) stall prior to completing it (**Figure 8C**) due to the inability or inefficiency in carrying out the subsequent dNMP incorporation steps (**Figure 8B**) (1). These results indicate that εA and T<>T lesions are very strong blocks to pol δ holoenzyme progression, particularly during insertion, suggesting that in lagging strand templates pol δ holoenzymes do not significantly contribute to the bypass of these lesions, if at all. Thus, DDT is likely required to execute complete bypass of these lesions in lagging strand templates.

### Pol δ holoenzymes contribute to the bypass of 3MeC, FapyG, Ab, Tg, and 8OxoG lesions in lagging strand templates

Human pol δ holoenzymes are markedly efficient at insertion opposite 3MeC, FapyG, Ab, Tg, and 8OxoG lesions (**Figure 8B**). However, bypass of each lesion is compromised by relatively inefficient extension (**Figure 8B**). This results in a distribution of pol δ holoenzyme fates upon encountering these lesions. Less than half of progressing pol δ holoenzymes stall upon encounter (15.8 ± 0.6 % - 51.8 ± 5.2%). The remainder either initiate bypass and then stall during extension (25.3 ± 0.7% - 58.5 ± 0.8%) or complete bypass (7.13 ± 0.32 % – 36.4 ± 1.1%) (**Figure 8C**). Collectively, this indicates that progressing pol δ holoenzymes contribute to the bypass of the majority (48.2 ± 5.2 % - 84.2 ± 0.6 %) of 3MeC, FapyG, Ab, Tg, and 8OxoG lesions they encounter, either by initiating bypass or executing complete bypass. Based on these results, we propose that these lesions are bypassed in lagging strand templates by both pol δ-independent and -dependent mechanisms (**Figure 9**). For pol δ-independent mechanisms (**Figure 9**, *Top*), pol δ holoenzymes stall upon encountering these lesions, activating DDT pathways that are ultimately responsible for the entirety of lesion bypass. *Ex vivo* and *in vivo* studies of human cells suggest that TLS is the primary DDT pathway involved in the bypass of these lesions (37,40,53-61) and numerous *in vitro* studies from independent labs have demonstrated complete bypass of these lesions by one or more human TLS polymerases (15,36,37,40,58,62-65).

Following lesion bypass, pol δ holoenzymes resume DNA synthesis downstream of the offending lesion.

In pol δ-dependent mechanisms, progressing pol δ holoenzymes may initiate lesion bypass and then stall during extension. Stalling elicits TLS polymerases to perform extension past these lesions, completing bypass, and then pol δ holoenzymes resume DNA synthesis during elongation. Many *in vitro* studies from independent labs demonstrated that several human TLS polymerases can extend correct and mismatched primer termini past each of these lesions (15,37,40,62,63,66). Based on our results (**Figure 8C**), this is the primary mechanism for pol δ-dependent bypass of these lesions, particularly 3MeC and Ab. Alternatively, progressing pol δ holoenzymes may execute complete bypass in the absence of TLS polymerases (**Figure 9**, *Bottom*) and continue progressing (i.e., elongating). Based on the results presented in **Figure 8C**, this is a secondary mechanism for pol δ-dependent bypass of these lesions but is notably significant for FapyG, Tg, and 8OxoG lesions. Previous investigations of human cells support pol δ-dependent mechanisms for lesion bypass. In particular, studies from the Livneh laboratory suggest that pol δ is involved in the bypass of 8OxoG lesions and is required for the bypass of Ab lesions (55,59). To the best of our knowledge, comparable cellular studies addressing the contributions of pol δ holoenzymes to the bypass of 3MeC, FapyG, or Tg lesions have yet to be reported. Such cellular studies are the focus of future efforts.

### Contribution of proofreading by pol δ holoenzymes during lesion bypass

Collectively, the results from our previous and present study indicate that proofreading by pol δ holoenzymes does not deter their progression at any lesion analyzed (**Figure 9**)(1). Rather, it significantly promotes progression at 7 of the 10 lesions (3MeC, Ab, 8OxoG, U, FapyG, Tg, and O6MeG). Interestingly, proofreading by pol δ holoenzymes significantly enhances insertion opposite 3MeC and Ab lesions (i.e., initiating bypass), while having a relatively smaller effect on the (complete) bypass of these lesions. Under the conditions of the assays in our investigations, lesions are encountered only by progressing pol δ holoenzymes (rather than pol δ alone) during initial binding events with P/T DNA substrates. To the best of our knowledge, comparable studies and analyses of human lagging strand replication have yet to be reported. Based on the assay design, the aforementioned results for the 10 lesions analyzed suggest that proofreading by pol δ holoenzymes promotes their progression at certain lesions by affording multiple attempts at one or more dNMP incorporation steps involved in lesion bypass.

## Supplementary Data Statement

Supplementary Data are available at

## Acknowledgements

We would like to thank all members of the Hedglin lab for their efforts in reviewing/proofreading the current manuscript.

## Author contributions

R.L.D. expressed, purified, and characterized all proteins. R.L.D., J.A.C., H.W., and S.G. performed the experiments and analyzed the data. M.H., R.L.D., S.G., and M.M.G. designed the experiments and analyzed the data. M.H., R.L.D. and M.M.G. wrote the paper.

