## Supporting Information for "A human high-fidelity DNA polymerase holoenzyme has a wide range of lesion bypass activities"

**Affiliations**:
[1]Department of Chemistry, The Pennsylvania State University, University Park, PA 16802
[2]Department of Chemistry, Johns Hopkins University, Baltimore, MD 21218

### Experimental Methods

*FRET-based assay to monitor extent of RFC-catalyzed loading of PCNA.* All experiments were performed at room temperature (23 ± 2 °C) in 1X Replication Buffer (25 mm HEPES, pH 7.5, 10 mm $Mg(OAc)_2$, 125 mm KOAc) supplemented with 1 mm DTT, and the final ionic strength was adjusted to physiological (200 mm) by the addition of appropriate amounts of KOAc. All measurements were done in 16.100F-Q-10/Z15 sub-micro fluorometer cells (Starna Cells) in a Horiba Scientific Duetta-Bio fluorescence/absorbance spectrometer. Excitation and emission slit widths are each set to 5 nm, unless indicated otherwise. Reaction solutions are excited at 514 nm and the fluorescence emission intensities ($I$) are simultaneously monitored at 563 nm ($I_{563}$, Cy3 FRET donor fluorescence emission maximum) and 665 nm ($I_{665}$, Cy5 FRET acceptor fluorescence emission maximum) over time, recording $I$ every 0.17 s. For each time point, $E_{FRET}$ is calculated where $E_{FRET} = \frac{I_{665}}{I_{665}+I_{563}}$. $E_{FRET}$ values for complete loading of PCNA onto a P/T junction were calculated based on a published FRET-based assay (1-10). A solution containing 110 nM of a Bio-Cy3P/T DNA (**Figure S2**), 440 nM neutravidin and 1 mM ATP is pre-incubated with RPA (330 nM heterotrimer). Then, Cy5-PCNA (100 nM homotrimer) is added, the resultant solution is transferred to a fluorometer cell, and the cell is placed in the instrument. $I_{665}$ and $I_{563}$ are monitored until both stabilize for at least 1 min. Within this stable region, $E_{FRET}$ values are calculated and averaged to obtain the $E_{FRET}$ value observed prior to addition of RFC•ATP complexes. Finally, pre-formed RFC•ATP complexes (100 nM RFC heteropentamer, 1 mM ATP) are added, the resultant solution is mixed via pipetting, and $E_{FRET}$ is monitored beginning 10 s after the addition of RFC•ATP complexes. Data is plotted as a function of time with time courses adjusted for the time between the addition of RFC and the recording of $E_{FRET}$ ($\Delta t$ = 10 s).

*Primer Extension Analysis for templates containing a T<>T or the corresponding native di-thymine sequence*. For a T<>T or the corresponding native di-thymine sequence that starts at dNMP incorporation *i*, bypass requires dNMP incorporation opposite the 3′ pyrimidine (i.e., insertion 1 at *i*), the 5′ pyrimidine (i.e., insertion 2 at *i* +1), and the 1$^{st}$ template nucleotide downstream of the 5′ pyrimidine (i.e., extension at $i + 2$). Accordingly, the probabilities of insertion 1, insertion 2, extension, and bypass are equal to $P_i$, $P_{i+1}$, $P_{i+2}$, and $P_i \times P_{i+1} \times P_{i+2}$, respectively. These probabilities are utilized to calculate the efficiencies of insertion 1, insertion 2, extension, and bypass for a T<>T, as described in the main text. Encounter of a CPDT<>T or the corresponding native di-thymine sequence that starts at dNMP incorporation step *i* is indicated by the presence of primer extension products resulting from the completion of dNMP incorporation steps up to and including *i*-1. These particular primer extension products are collectively referred to as encounter products. The fraction of progressing pol δ holoenzymes that encounter a T<>T or the corresponding native di-thymine sequence that starts at dNMP incorporation step *i* and subsequently stall (via dissociation of pol δ) at dNMP incorporation step *i*-1 (i.e., during insertion) or any dNMP incorporation step downstream is defined as the band intensity of the aborted primer extension product resulting from completion of that particular dNMP incorporation step divided by the sum of the band intensities of all encounter products. These values are utilized to determine the distribution (%) of events that occur after progressing pol δ holoenzymes encounter a T<>T or the corresponding native di-thymine sequence that starts at dNMP incorporation step *i*. Specifically, the events are 1) stall during encounter (i.e., stall during insertion 1); 2) initiate bypass then stall (i.e., complete insertion then stall during insertion 2 or extension) and; 3) complete bypass (i.e., complete insertion 1, insertion 2, and extension).

### Supporting Figures

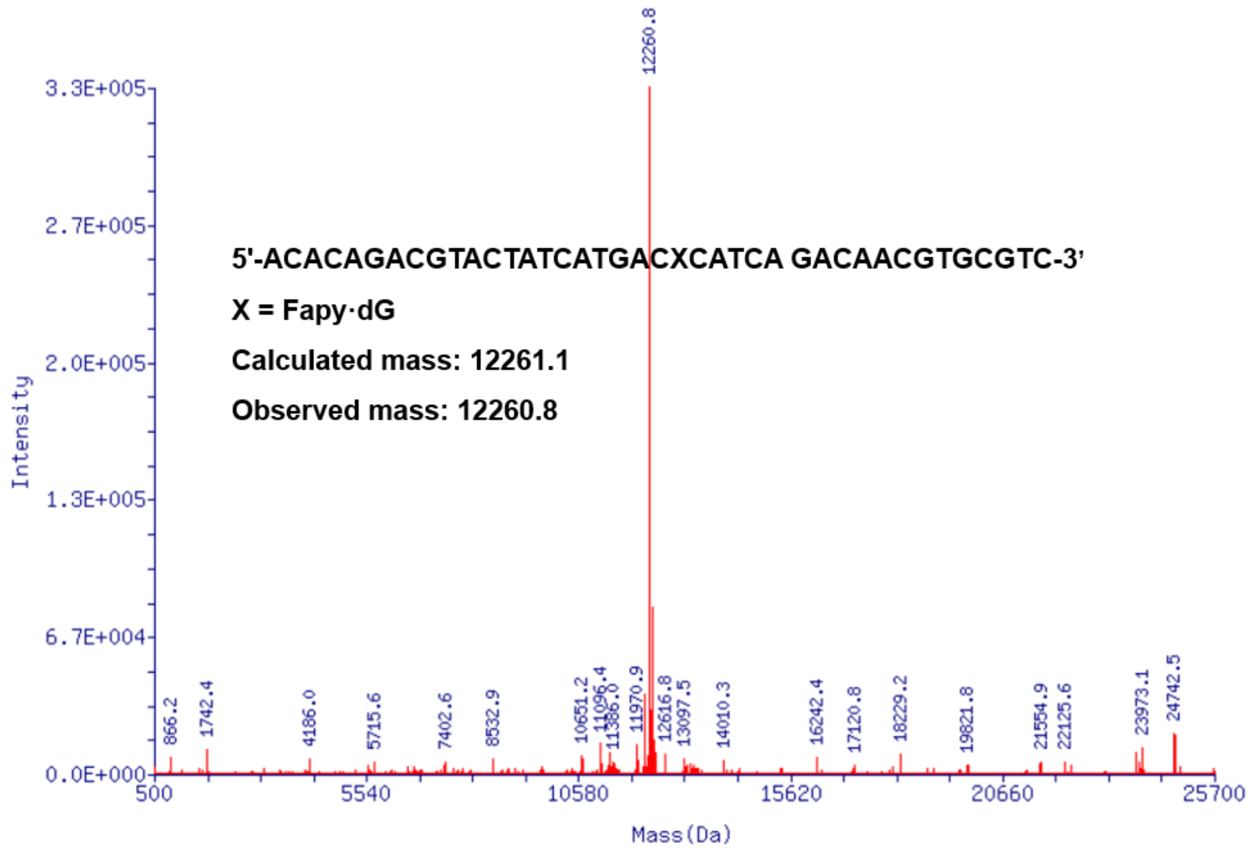


**Figure S1**. Deconvoluted mass spectrum of the 40-mer FapyG-containing oligonucleotide.

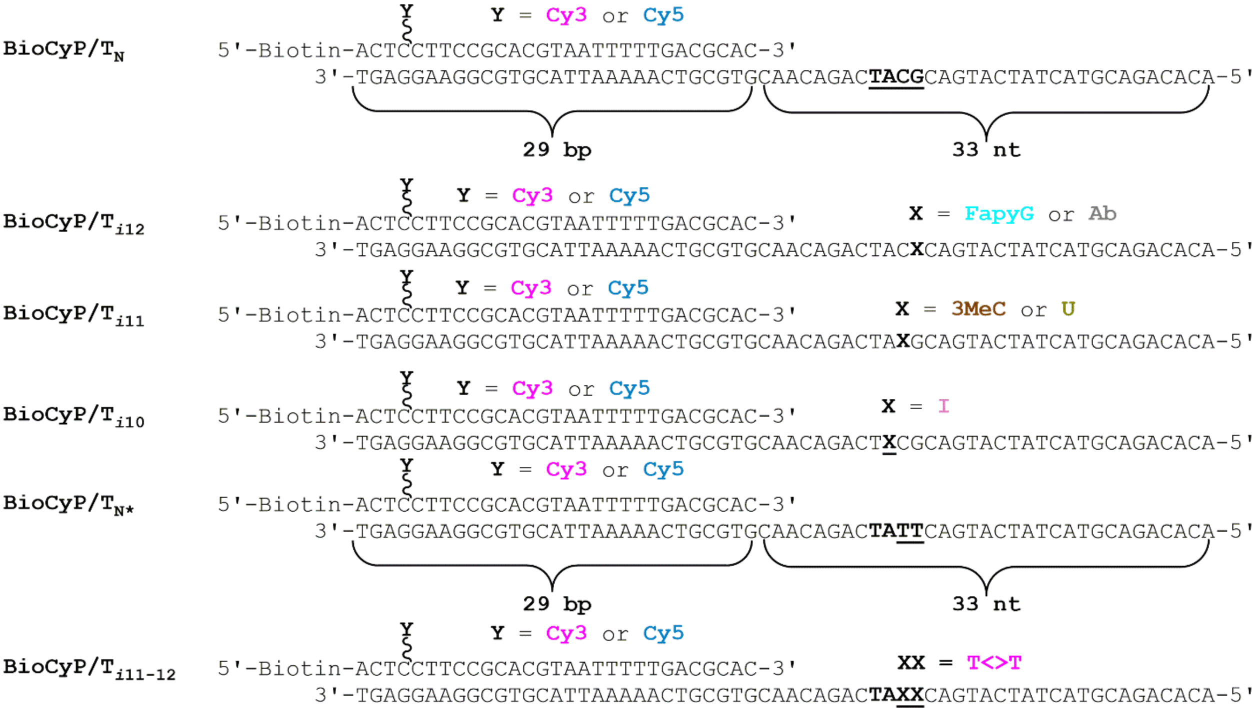


**Figure S2**. DNA substrates utilized in the current study. For all P/T DNA substrates, the sequences and lengths (29 bp) of the double strand DNA (dsDNA) regions are identical and in agreement with the requirements for assembly of a single PCNA ring onto DNA by RFC (1,4,7). The lengths of the ssDNA regions adjacent to the 3′-end of the P/T junctions are identical (33 nt) and accommodate 1 RPA molecule (11-13). RPA prevents loaded PCNA from sliding off the ssDNA end of the substrate (4). When pre-bound to neutravidin, the biotin attached to the 5’ end of a primer strand prevents loaded PCNA from sliding off the dsDNA end of the substrate. The underlined DNA sequence in the undamaged, control P/T DNA substrates (BioCyP/$T_N$ and BioCyP/$T_{N*}$) highlights the native template nucleobases that are chemically modified (i.e., damaged) to DNA lesions.

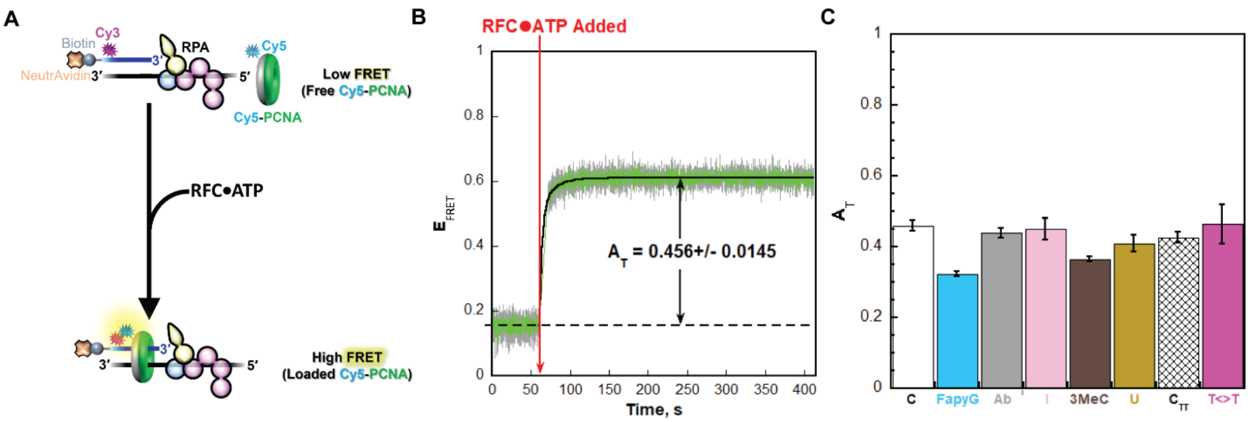


**Figure S3.** RFC-catalyzed loading of PCNA onto DNA. (**A**) Schematic representation of the assay. The DNA substrates are P/T DNA substrates (**Figure S2**) in which the primer contains an internal Cy3 FRET donor (Y = Cy3, **Figure S2**) 4 nt from its 5′ terminus. The "back face" of PCNA (shown in grey) is labeled with a Cy5 FRET acceptor. Cy5-PCNA is loaded onto the DNA substrate by the human clamp loader, RFC, such that the Cy5 FRET donor on the "back face" of PCNA faces the Cy3 FRET donor near the 5′ terminus of the primer strand and the "front face" of PCNA (shown in green) is oriented towards the P/T junction where DNA synthesis emanates from. Cy5-PCNA is loaded by RFC in the presence of excess RPA and $E_{FRET}$ is monitored over time. Under these conditions, all Cy5-PCNA is loaded onto a native (i.e., undamaged) BioCyP/T DNA substrate (**Figure S2**, either BioCy3P/$T_N$ or BioCy3P/$T_{N^*}$ ) by RFC and stabilized by RPA and the biotin/neutravidin blocks that prevent diffusion of PCNA off of the DNA (1-9). (**B**) FRET data observed for the BioCy3P/$T_N$ DNA substrate. The $E_{FRET}$ trace is the average of at least three independent traces and the standard error is shown in grey. The time at which RFC•ATP complexes are added is indicated by a red arrow. The portion of the $E_{FRET}$ trace observed prior to the addition of RFC•ATP complexes represents the complete absence of interactions between the BioCy3P/$T_N$•RPA complexes and Cy5-PCNA and is fit to a flat line that is extrapolated to the axis limits. The portion of the $E_{FRET}$ trace observed after the addition of RFC•ATP complexes is fit to a double exponential rise and the sum of the amplitudes for the two phases (overall, total amplitude, $A_T$) is reported in the graph. $A_T$ indicates the change in $E_{FRET}$ observed when Cy5-PCNA is stably loaded onto a BioCy3P/T DNA substrate (**Figure S2**). Thus, $A_T$ directly reports on the extent of stable assembly of PCNA onto DNA. (**C**) $A_T$ data. Each column represents the average $\pm$ SEM of three independent experiments for a respective BioCyP/T DNA substrate. Data for the native control BioCy3P/$T_N$ (Control, "C") serves as a reference for the data observed for the BioCy3P/$T_{i12}$ with either FapyG, O6MeG or an abasic site mimic at dNTP incorporation step *i* = 12 , BioCy3P/$T_{i11}$ with an I at *i* = 11, and the BioCy3P/$T_{i10}$ with either a 3MeC or a U at *i* = 10. Data for the the native control BioCy3P/$T_{N^*}$ (Control TT, "$C_{TT}$") serves as a reference for the data observed for the BioCy3P/$T_{i11\text{-}12}$ with a T<>T at *i* = 11 – 12. The $A_T$ value observed for the native control DNA substrates (BioCy3P/$T_N$ and BioCy3P/$T_{N^*}$) agrees with that observed previously (1-9), indicating that all Cy5-PCNA is loaded onto the P/T junctions and stabilized. Significant $A_T$ values are observed for all P/T DNA substrates containing a DNA lesion at least 9 nt downstream of the P/T junction and the observed $A_T$ values are in good agreement with the values observed for their respective reference control DNA substrate. Altogether, this indicates that PCNA is stably loaded onto damaged P/T DNA substrates and DNA lesions do not significantly affect the stable assembly of PCNA onto a P/T junction upstream of the lesion.
